# First report and multilocus genotyping of *Enterocytozoon bieneusi* from Tibetan pigs in southwestern China

**DOI:** 10.1101/327767

**Authors:** Run Luo, Leiqiong Xiang, Haifeng Liu, Zhijun Zhong, Li Liu, Lei Deng, Yuan Song, Ling Liu, Xiangming Huang, Ziyao Zhou, Hualin Fu, Yan Luo, Guangneng Peng

**Affiliations:** The Key Laboratory of Animal Disease and Human Health of Sichuan Province, College of Veterinary Medicine, Sichuan Agricultural University, Chengdu, Sichuan Province 611130, China; Chengdu Giant Panda Breeding Research Base, Chengdu, Sichuan Province 625001, China

## Abstract

*Enterocytozoon bieneusi* is a common intestinal pathogen and a major cause of diarrhea and enteric diseases in a variety of animals. While the *E. bieneusi* genotype has become better-known, there are few reports on its prevalence in the Tibetan pig. This study investigated the prevalence, genetic diversity, and zoonotic potential of *E. bieneusi* in the Tibetan pig in southwestern China. Tibetan pig feces (266 samples) were collected from three sites in the southwest of China. Feces were subjected to PCR amplification of the internal transcribed spacer (ITS) region. *E. bieneusi* was detected in 83 (31.2%) of Tibetan pigs from the three different sites, with 25.4% in Kangding, 56% in Yaan and 26.7% in Qionglai. Age group demonstrated the prevalence of *E. bieneusi* range from 24.4%(aged 0 to 1 years) to 44.4%(aged 1 to 2 years). Four genotypes of *E. bieneusi* were identified: two known genotypes EbpC (n=58), Henan-IV (n=24) and two novel genotypes, SCT01 and SCT02 (one of each). Phylogenetic analysis showed these four genotypes clustered to group 1 with zoonotic potential. Multilocus sequence typing (MLST) analysis three microsatellites (MS1, MS3, MS7) and one minisatellite (MS4) revealed 47, 48, 23 and 47 positive specimens were successfully sequenced, and identified ten, ten, five and five genotypes at four loci, respectively. This study indicates the potential danger of *E. bieneusi* to Tibetan pigs in southwestern China, and offers basic data for preventing and controlling infections.

## Introduction

Microsporidia are obligate intracellular eukaryotic pathogens, classified as fungi, which are composed of approximately 1300 species in 160 genera[1]. To date, 17 microsporidia species are known to infect humans, and of these, *E. bieneusi* is the most prevalent, accounting for over 90% of cases of human microsporidiosis[2]. Since its first detection in an HIV/AIDS patient in 1985, a growing literature attests to *E. bieneusi* expanding range of hosts [3–5]. In humans, infection by microsporidia results in self-limiting diarrhea and malabsorption, most seriously, immunocompromised and immunocompetent patients more susceptible to *E. bieneusi* infection[6]. Normally, fecal-oral routes serve as the main infection pathways in humans and animals, while human inhalation of *E. bieneusi* spores has also been documented[7, 8].

PCR-based molecular techniques may be used to analyze the *E. bieneusi* genome, and for diagnosis. Based on the nested PCR amplification of internal transcribed spacers (ITS) of small subunits of ribosomal rRNA (SSU rRNA), over 240 *E. bieneusi* genotypes have been identified globally[9–11]. Phylogenetic analysis reveals that these genotypes clustered into nine groups. Group 1 is considered zoonotic, and is composed of genotypes from humans and a few animals, while groups 2–9 have particular host associations or are found in wastewater[5, 11]. To better comprehend *E. bieneusi* genetic diversity and molecular characteristics, high-resolution multi-locus sequence typing (MLST) using three microsatellites (MS1, MS3 and MS7) and one minisatellite (MS4) as markers was used to explore genotype taxonomy and transmission routes [9, 12, 13].

In the southwest of China, Tibetan pigs are widely kept for livelihood and are economically important for farmers, especially on the plateau. Tibetan pigs have firm black hair which differs from that of the common pig, and are sturdy, outdoor foragers. They may act as reservoirs for *E. bieneusi* spores and zoonotic transmission of disease. Although much research has been carried out on *E. bieneusi*[14–16], few studies have examined its epidemiology or Tibetan pig-associated genomes in China[17, 18], and Tibetan pigs in southwestern China have been entirely unstudied. Therefore, this study aimed to establish the incidence and molecular characteristics of *E. bieneusi* in Tibetan pigs, to use ITS and MLST to evaluate its genetic diversity, and to assess the potential for zoonotic transmission of microsporidiosis between Tibetan pigs and humans.

## Materials and methods

### Ethics statement

The study was conducted in accordance with the Research Ethics Committee and the Animal Ethics Committee of Sichuan Agricultural University. Prior to fecal specimen collection, permission was obtained from the keepers of the animals whenever possible.

### Collection of Tibetan pig fecal specimens

Fresh fecal specimens were collected from 266 Tibetan pigs during June-October 2017. Samples were obtained mainly from three cities in Sichuan province, southwestern of China, including Yaan (n=50), Kangding (n=201) and Qionglai (n=15) (Table 1). The ages of Tibetan pigs sampled ranged from 1 to 2 years. Specimens were collected using sterile disposable latex gloves immediately after being defecated on to the ground, and transferred into 50 ml plastic containers. All experimental Tibetan pigs were not any diarrheic or gastrointestinal conditions. Samples were stored at 4 °C in 2.5% (w/v) potassium dichromate.

**Table 1.**
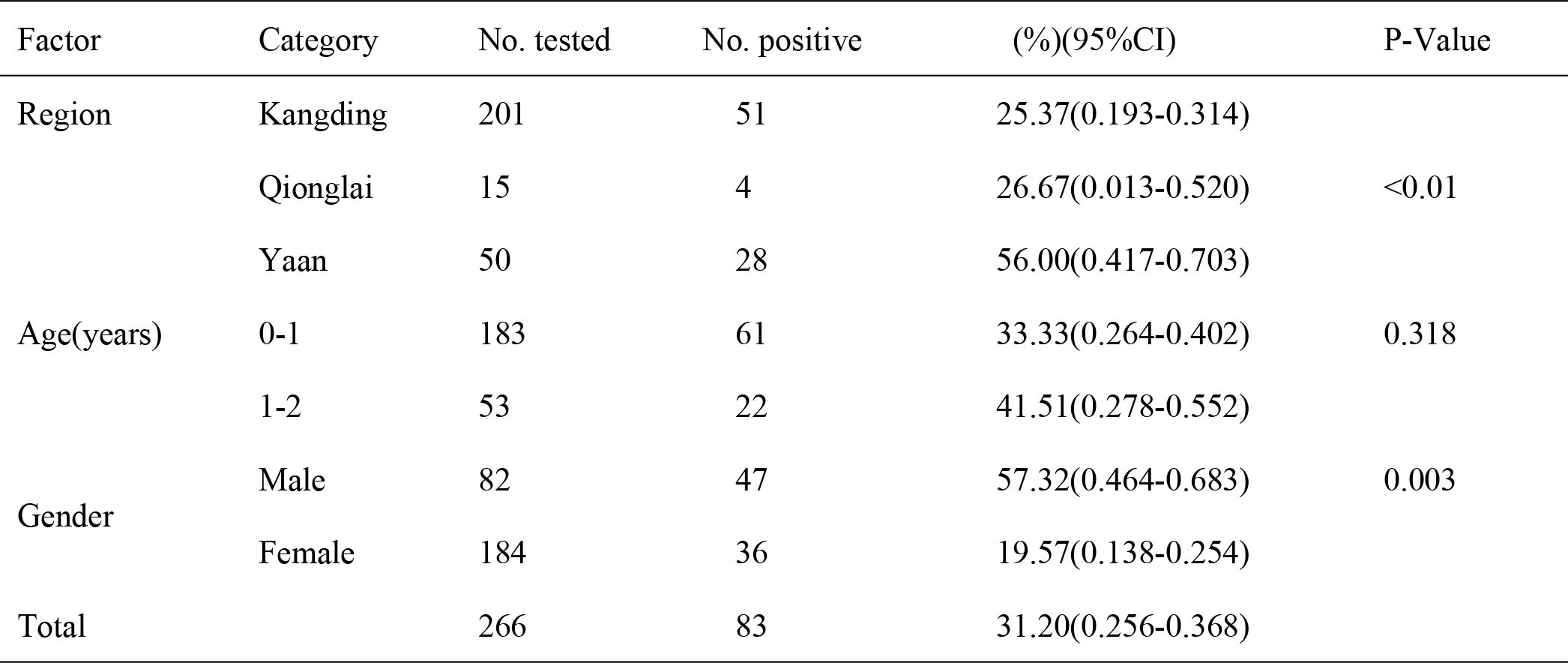
Factors associated with prevalence of *Enterocytozoon bieneusi* in Tibetan pigs in southwestern China

| Factor | Category | No. tested | No. positive | (%)(95%CI) | P-Value |
| --- | --- | --- | --- | --- | --- |
| Region | Kangding | 201 | 51 | 25.37(0.193-0.314) | <0.01 |
|  | Qionglai | 15 | 4 | 26.67(0.013-0.520) |  |
|  | Yaan | 50 | 28 | 56.00(0.417-0.703) |  |
| Age(years) | 0-1 | 183 | 61 | 33.33(0.264-0.402) | 0.318 |
|  | 1-2 | 53 | 22 | 41.51(0.278-0.552) |  |
| Gender | Male | 82 | 47 | 57.32(0.464-0.683) | 0.003 |
|  | Female | 184 | 36 | 19.57(0.138-0.254) |  |
| Total |  | 266 | 83 | 31.20(0.256-0.368) |  |

### DNA extraction

Before conducting DNA extraction, potassium dichromate was removed from the fecal samples with distilled water by centrifugation for 10 minutes at 1500 × g, three times. Genomic DNA was extracted from 200 mg of washed fecal matter using the EZNA1 Stool DNA kit (Omega Biotek, Norcross, GA, USA). Prior to use in PCR analysis, DNA was stored and frozen at −20 °C.

### PCR amplification

*E. bieneusi* species/genotypes were determined using a nested PCR amplification of the entire ITS region, and positive specimens were further detected by MLST analyses using the MS1, MS3, MS4, and MS7 loci. The primers and cycling parameters implemented for these reactions were as previously described[12, 19]. Negative controls were included in all PCR analyses. The secondary PCR products were subjected to electrophoresis in a 1.5% agarose gel and visualized under UV light by staining the gel with GoldView (Solarbio, China).

### Nucleotide sequencing and phylogenetic analysis

Secondary PCR amplicons of anticipated size were sequenced in both directions by Life Technologies (Guangzhou, China) with an ABI 3730DNA Analyzer (Applied Biosystems, Foster City, CA, USA) using the BigDye^®^ Terminator v3.1 cycle sequencing kit. Sequence accuracy was confirmed by bidirectional sequencing, and new PCR secondary products were re-sequenced, if necessary. To identify the *E. bieneusi* genotype, the sequences generated were respectively aligned with known reference sequences using BLAST and ClustalX 1.83. Mega 7.0 was used to construct the phylogenetic tree using the neighbor-joining (NJ) method (the Kimura two parameter model) with 1000 bootstrap replicates[20]. Novel genotype(s) of *E. bieneusi* were named according to the established system of nomenclature[21].

### Statistical analysis

The variation in *E. bieneusi* infection rates in Tibetan pigs between different areas, gender, and ages were compared using the Chi-square test. All tests were two-sided, with P <0.05 indicating statistical significance. SPSS version 22.0 was used on all data. 95% confidence intervals (95% CIs) were calculated to explore the strength of the association between *E. bieneusi* occurrence and each factor.

### Nucleotide sequence accession numbers

Representative nucleotide sequences of *E. bieneusi* isolates were deposited in GenBank under accession numbers from MG581429-MG581432 for ITS sequences and MH142189-MH142213 for the microsatellite (MS1, MS3, and MS7) and minisatellite (MS4) loci.

## Results

### Occurrence of *E. bieneusi* in Tibetan pigs

Of the 266 Tibetan pigs sampled 83 (31.3%) were PCR-positive for *E. bieneusi*. The epidemiology and genotypes of *E. bieneusi* in different areas are given in Table 2. Infection rates detected in Tibetan pigs were 25.4%, 56% and 26.6% in Kang ding, Ya an and Qiong lai, respectively. Infection rates by age and gender are given in Table 1. Differences between the three areas were significant (χ^2^=17.648, df =2, p<0.01). In addition, the female Tibetan pig groups (17.7%, 47/266,) had a higher *E. bieneusi* prevalence than the male groups (13.5%, 36/266,). The difference in infection rate was also significant (χ^2^=8.906, df =1,P=0.003); however, in the present study, high infection rates were observed in 1–2 year-old pigs (41.51%, 22/53) and 0–1 year-olds (33.33%, 61/183); however, these rates were not significantly different (χ^2^ = 1.240, df =1, P>0.05).

**Table 2.** Occurrence and genotypes of *E. bieneusi* in Tibetan pigs from different cities in southwest China.

| Region | Farm ID | Prevalence (%) | Genotypes (n) |
| --- | --- | --- | --- |
| Kangding | Farm 1 | 31/102(30.40) | EbpC(18),Henan-IV(n=13) |
|  | Farm 2 | 20/99(20.20) | EbpC(12),Henan-IV(n=8) |
| Yaan | Farm 3 | 14/28(50.00) | EbpC(n=14) |
|  | Farm 4 | 10/22(45.45) | EbpC(n=12),SCT01(n=1),SCT02(n=1) |
| Qionglai | Farm 5 | 4/15(26.67) | EbpC(n=4) |
| Total |  | 83/266(31.20) | EbpC(58),Henan-IV(n=23),SCT01(n=1),SCT02(n=1) |

### Genotype distribution of *E. bieneusi* in Tibetan pigs

Nucleotide sequences from ITS-PCR were obtained from the 83 *E. bieneusi*-positive specimens. Four genotypes were detected, including two known genotypes (EbpC, Henan-IV) and two novel genotypes, which were named SCT01 and SCT02 (Table2). Genotype EbpC was the most prevalent (21.8%, 58/266), and was detected in samples from all three cities. Genotype Henan-IV was only found in Kang ding (8.6%, 23/266). The novel genotypes SCT01 (0.3%, 1/266) and SCT02 (0.3%, 1/266) were only found in single specimens, both of which came from Ya an, and are the first newly-detected *E. bieneusi* genotypes from Tibetan pigs.

### Phylogenetic relationships of *E. bieneusi* ITS genotypes

Phylogenetic analysis based on ITS gene sequences, the four *E. bieneusi* genotypes obtained from the present study (two known and two novel genotypes) were classed as a single group (group 1) and further clustered into subgroup 1d, indicating zoonotic potential (Fig 1).

**Fig 1.**
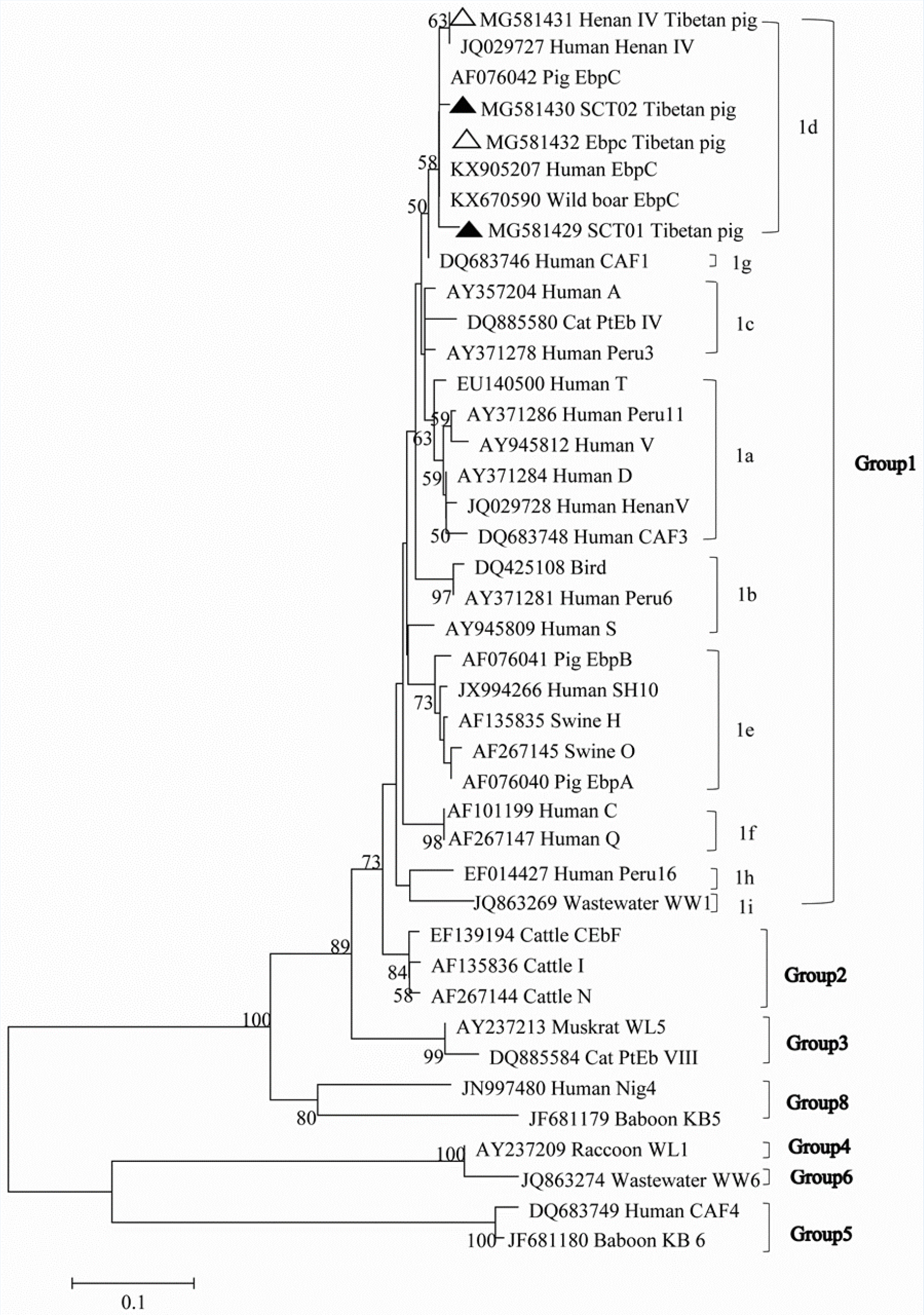
Phylogenetic relationship of *Enterocytozoon bieneusi* groups, the relationship between *E. bieneusi* genotypes identified in this study and other known genotypes deposited in the GenBank was inferred by a neighbor-joining analysis of ITS sequences based on genetic distance by the Kimura-2-parameter model. The numbers on the branches represent percent bootstrapping values from 1,000 replicates, with more than 50% shown in tree. Each sequence is identified by its accession number, genotype designation, and host origin. Genotypes with black triangles and open triangle are novel and known genotypes identified in this study, respectively.

### Multilocus genotyping of *E. bieneusi*

Positive specimens were further characterized by PCR analyses of MS4, MS1, MS3 and MS7 to improve taxonomy and population genotypes of *E. bieneusi*. 47, 48, 23 and 47 *E. bieneusi* isolates were amplified at the MS1, MS3, MS4, and MS7 loci, respectively, but only 12 samples were PCR-positive simultaneously at all four loci. Four distinct MLGs were observed in Henan-IV and six distinct MLGs in EbpC, named MLG1-4 and MLG5-10, respectively (Table 3). These results reveal high genetic diversity in the Henan-IV and EbpC genotypes of *E. bieneusi* in Tibetan pigs. Table3 Multilocus characterization of *Enterocytozoon bieneusi* isolates in Tibetan pigs in Southwestern China

**Table 3.**

| Genotype (Synonym) | Host | Location | Isolate | Reference |
| --- | --- | --- | --- | --- |
| EbpC (E, Peru4, WL13, WL17) | Pig | Shanghai | 3 | [12] |
|  | Pig | Heilongjiang | 10 | [12] |
|  | Pig | Heilongjiang | 3 | [31] |
|  | Pig | Heilongjiang | 3 | [32] |
|  | Pig | Jilin | 1 | [33] |
|  | Pig | Mongolia | 1 | [33] |
|  | Tibetan pig | Sichuan | 58 | This study |
|  | Red panda | Shanxi | 5 | [34] |
|  | Human | Shanghai | 1 | [35] |
|  | Human | Henan | 39 | [29] |
|  | Human | Heilongjiang | 11 | [36] |
|  | Human,pig,Monkey | Guangxi | 4 | [37] |
|  | Squirrel | Sichuan | 3 | [25] |
|  | Wild boar | Sichuan | 85 | [30] |
|  | nonhuman primates | Hebei | 1 | [38] |
|  | nonhuman primates | Hubei | 3 | [38] |
|  | nonhuman primates | Hunan | 4 | [38] |
|  | nonhuman primates | Being | 2 | [38] |

## Discussion

In the present study, infection rates of varied between 25.4%–56% in different districts, with an overall infection rate of 31.2%. This rate is lower than the documented prevalence of *E. bieneusi* for: wild boars in Sichuan province, China (41.2%), domestic pigs in Jilin province, China (45.1%), wild boars in central Europe (33.3%) and pigs in the State of Rio de Janeiro, Brazil (59.3%) ^[14, 15, 22, 23]^. However, infection rates recorded in this study were higher than those for pigs in Jilin, China (16.4%), central Thailand (28.1%) and Japan (30%)[14, 16, 22]. Differences in infection rates between these studies may be largely attributable to climate and farming mode. Prevalences also varied across sample sites. Kang ding, the only site on the Western Sichuan Plateau, had a prevalence of 25.4%, possibly reflecting the area’s high temperatures, and UV radiation, which may limit survival of *E. bieneusi* spores and reduce transmission. Other factors influencing infection levels may include geo-ecological conditions, feeding/herd densities, herd management, sample size, and the condition of host animals. Differences in prevalence in Tibetan pigs between Ya an and Kang ding are thought to reflect differences between traditional and modern herd management and breeding technologies.

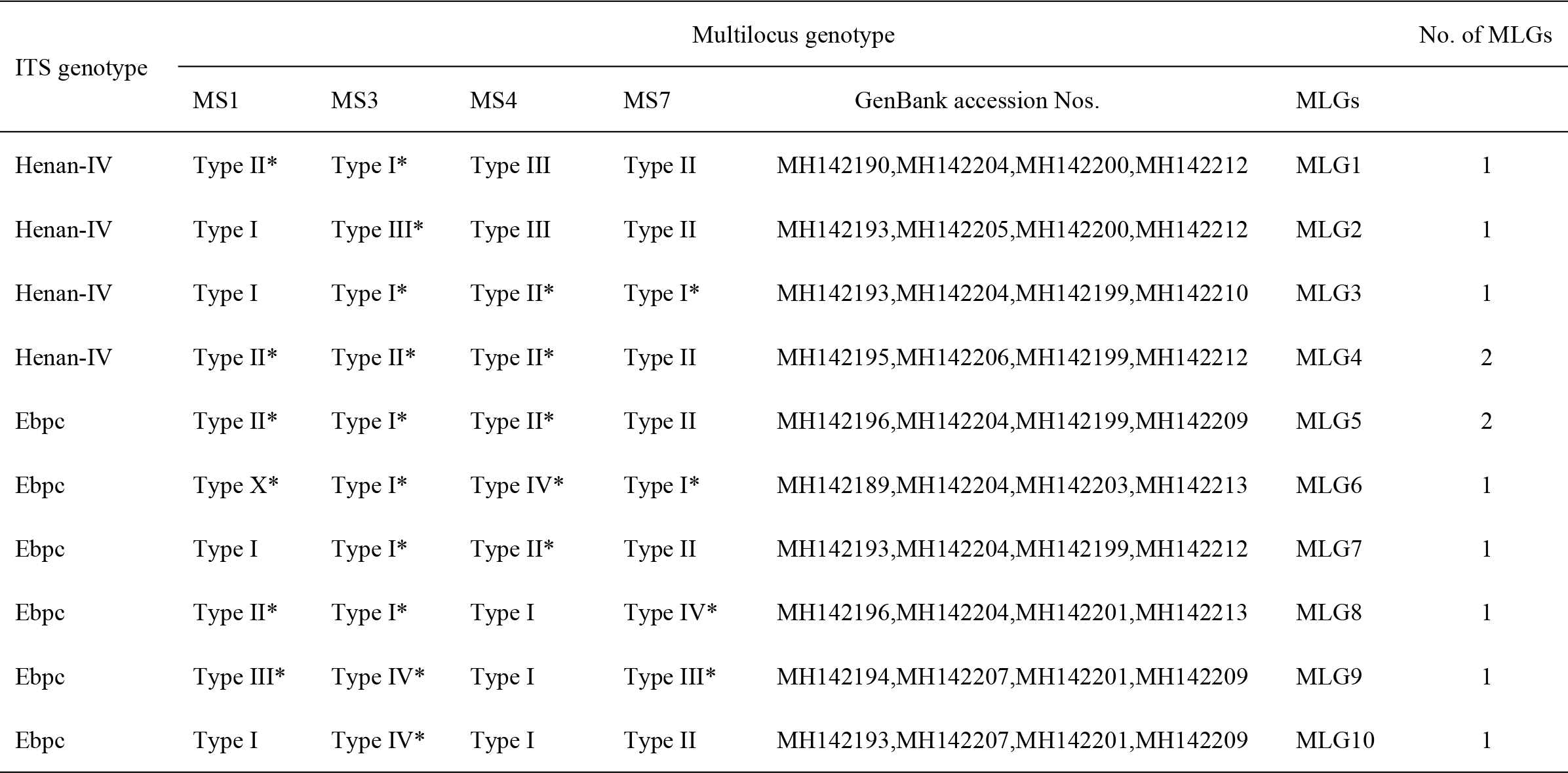

Of the four genotypes identified in this study EbpC was the most prevalent (21.8%, 58/266), and has been found in a number of animals, including cattle, dogs, cats, birds, non-human primates, bears, squirrels, sheep, foxes, deer, and humans[3, 5, 13, 23–28]. EbpC is the prevalent *E. bieneusi* genotype associated with pig infection in China, reflecting *E. bieneusi*’s dominance as a porcine parasite. In addition, we also detected 26 records of Henan-IV (solely in Ya an), a zoonotic genotype associated with human infections in Henan province in China, and thus far only recorded from China, where it demonstrates strict host specificity[29], occurring only in pigs and humans. To the best our knowledge, the two genotypes EbpC and Henan-IV, which were examined for the first time in Tibetan pigs in the present study, which may be a key reservoir host of these genotypes (Table 4).

**Table 4.**
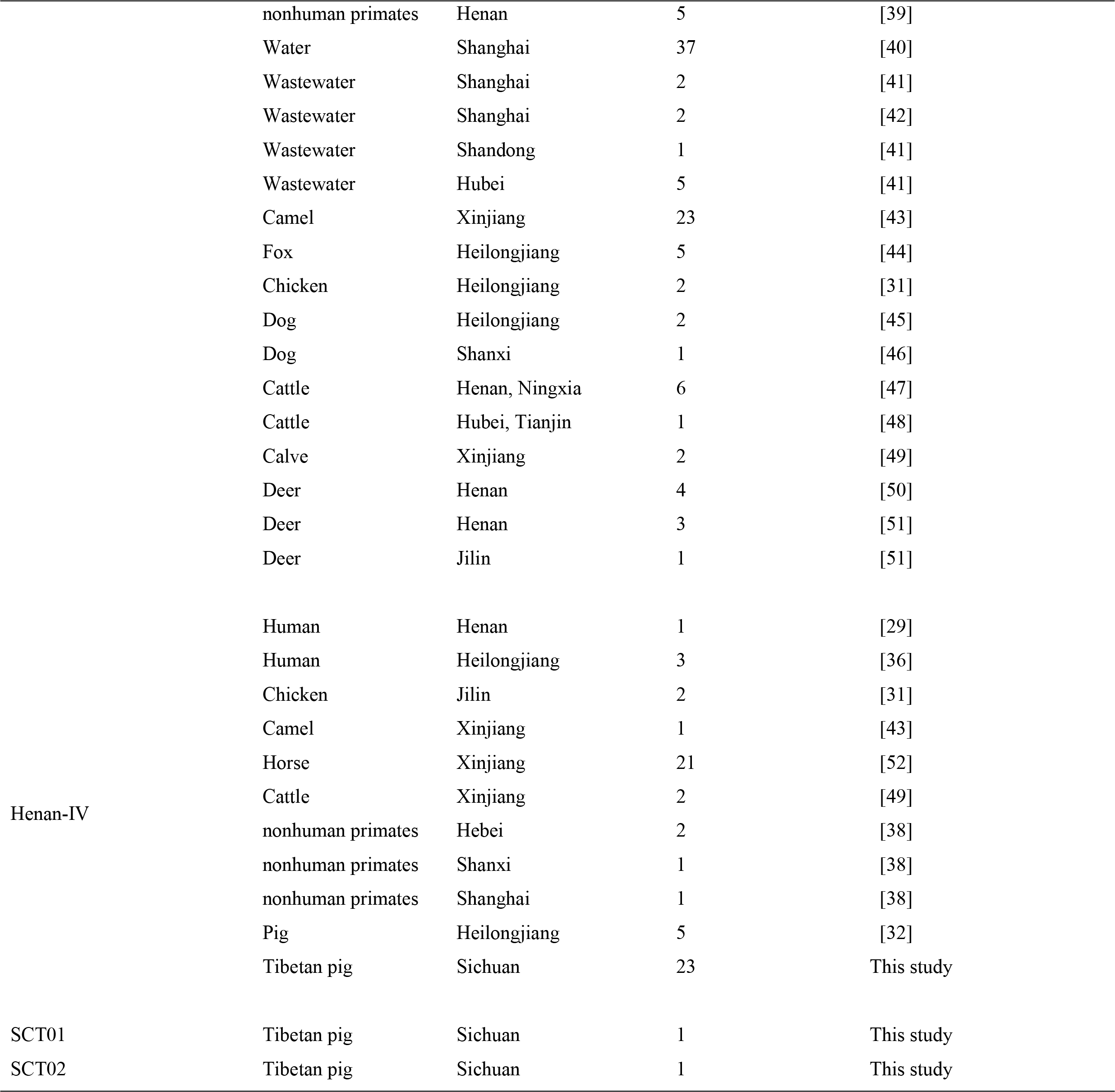
Host ranges and geographical distribution of *Enterocytozoon bieneusi* genotype in this study in China

|  |  |  |  |  |
| --- | --- | --- | --- | --- |
|  | nonhuman primates | Henan | 5 | [39] |
|  | Water | Shanghai | 37 | [40] |
|  | Wastewater | Shanghai | 2 | [41] |
|  | Wastewater | Shanghai | 2 | [42] |
|  | Wastewater | Shandong | 1 | [41] |
|  | Wastewater | Hubei | 5 | [41] |
|  | Camel | Xinjiang | 23 | [43] |
|  | Fox | Heilongjiang | 5 | [44] |
|  | Chicken | Heilongjiang | 2 | [31] |
|  | Dog | Heilongjiang | 2 | [45] |
|  | Dog | Shanxi | 1 | [46] |
|  | Cattle | Henan, Ningxia | 6 | [47] |
|  | Cattle | Hubei, Tianjin | 1 | [48] |
|  | Calve | Xinjiang | 2 | [49] |
|  | Deer | Henan | 4 | [50] |
|  | Deer | Henan | 3 | [51] |
|  | Deer | Jilin | 1 | [51] |
| Henan-IV | Human | Henan | 1 | [29] |
|  | Human | Heilongjiang | 3 | [36] |
|  | Chicken | Jilin | 2 | [31] |
|  | Camel | Xinjiang | 1 | [43] |
|  | Horse | Xinjiang | 21 | [52] |
|  | Cattle | Xinjiang | 2 | [49] |
|  | nonhuman primates | Hebei | 2 | [38] |
|  | nonhuman primates | Shanxi | 1 | [38] |
|  | nonhuman primates | Shanghai | 1 | [38] |
|  | Pig | Heilongjiang | 5 | [32] |
|  | Tibetan pig | Sichuan | 23 | This study |
| SCT01 | Tibetan pig | Sichuan | 1 | This study |
| SCT02 | Tibetan pig | Sichuan | 1 | This study |

ITS gene sequence analysis revealed two novel genotypes, SCT01 (n=1) and SCT02 (n=1), both of which were detected in Ya an and clustered into group 1 zoonotic genotypes with public health significance. Other genotypes in this group include Henan-III in humans and EbpC from humans or wild boars [13, 29, 30]. Modes of transmission and zoonotic potential of *E. bieneusi* genotypes remain poorly known, and further molecular epidemiology studies are required. MLST holds promise for ongoing investigation of *E. bieneusi* taxonomy and genetic diversity[12].

Nine, five, three and four novel genotypes were detected at MS1, MS3, MS4 and MS7 loci, respectively. Analysis of 12 samples at four gene loci identified eight novel MLGs, including three genotype EbpC MLGs and five genotype Henan-IV MLGs (Table 3).

## Conclusions

This study revealed an average *E. bieneusi* infection rate of 31.2% in three cities in Sichuan province, and is the first report of EbpC and Henan-IV in Tibetan pigs in China. Genetic diversity was characterized using MLST, and ten MLGs were identified. These results identify Tibetan pigs as possible vectors for zoonotic transmission of human microsporidiosis; Tibetan pigs widespread use and frequency of human contact make them a significant public health risk in southwest China. Thus, measures are needed to control the transmission of *E. bieneusi* and to develop effective vaccines and drugs for use in the event of widespread human microsporidiosis.

## Reference

1. Didier ES, Weiss LM: Microsporidiosis: current status. Current Opinion in Infectious Diseases 2006, 19(5):485.

2. Didier ES, Weiss LM: Microsporidiosis: not just in AIDS patients. Current Opinion in Infectious Diseases 2011, 24(5):490–495.

3. Wu J, Han JQ, Shi LQ, Zou Y, Li Z, Yang JF, Huang CQ, Zou FC: Prevalence, genotypes, and risk factors of Enterocytozoon bieneusi in Asiatic black bear (Ursus thibetanus) in Yunnan Province, Southwestern China. Parasitology Research 2018(6):1–7.

4. Zhang Q, Cai J, Li P, Wang L, Guo Y, Li C, Lei M, Feng Y, Xiao L: Enterocytozoon bieneusi genotypes in Tibetan sheep and yaks. Parasitology Research 2018, 117(1–2):1–7.

5. Zhang XX, Cong W, Lou ZL, Ma JG, Zheng WB, Yao QX, Zhao Q, Zhu XQ: Prevalence, risk factors and multilocus genotyping of Enterocytozoon bieneusi in farmed foxes (Vulpes lagopus), Northern China. Parasites & Vectors 2016, 9(1):1–7.

6. Santã-N M, Fayer R: Microsporidiosis: Enterocytozoon bieneusi in domesticated and wild animals. Research in Veterinary Science 2011, 90(3):363–371.

7. Zhao GH, Du SZ, Wang HB, Hu XF, Deng MJ, Yu SK, Zhang LX, Zhu XQ: First report of zoonotic Cryptosporidium spp., Giardia intestinalis and Enterocytozoon bieneusi in golden takins (Budorcas taxicolor bedfordi). Infection Genetics & Evolution 2015, 34:394–401.

8. Yan Z, Koehler AV, Tao W, Haydon SR, Gasser RB: First detection and genetic characterisation of Enterocytozoon bieneusi in wild deer in Melbourne’s water catchments in Australia. Parasites & Vectors 2018, 11(1):2.

9. Zhong Z, Tian Y, Song Y, Deng L, Li J, Ren Z, Ma X, Gu X, He C, Geng Y: Molecular characterization and multi-locus genotypes of Enterocytozoon bieneusi from captive red kangaroos (Macropus Rfus) in Jiangsu province, China. Plos One 2017, 12(8):e0183249.

10. Zhao W, Wang J, Yang Z, Liu A: Dominance of the Enterocytozoon bieneusi genotype BEB6 in red deer (Cervus elaphus) and Siberian roe deer (Capreolus pygargus) in China and a brief literature review. Parasite-journal De La Societe Francaise De Parasitologie 2017, 24(5):54.

11. Deng L, Li W, Zhong Z, Gong C, Liu X, Huang X, Xiao L, Zhao R, Wang W, Feng F: Molecular characterization and multilocus genotypes of Enterocytozoon bieneusi among horses in southwestern China. Parasites & Vectors 2016, 9(1):561.

12. Feng Y, Li N, Dearen T, Lobo ML, Matos O, Cama V, Xiao L: Development of a multilocus sequence typing tool for high-resolution genotyping of Enterocytozoon bieneusi. Applied & Environmental Microbiology 2011, 77(14):4822–4828.

13. Zhong Z, Li W, Deng L, Song Y, Wu K, Tian Y, Huang X, Hu Y, Fu H, Geng Y: Multilocus genotyping of Enterocytozoon bieneusi derived from nonhuman primates in southwest China. Plos One 2017, 12(5):e0176926.

14. Němejc K, Sak B, Květoňová D, Hanzal V, Janiszewski P, Forejtek P, Rajský, D, Kotková M, Ravaszová P, Mcevoy J: Prevalence and diversity of Encephalitozoon spp. and Enterocytozoon bieneusi in wild boars (Sus scrofa)in Central Europe. Parasitology Research 2014, 113(2):761.

15. Fiuza VRS, Oliveira FCR, Fayer R, Santín M: First report of Enterocytozoonbieneusi in pigs in Brazil. Parasitology International 2015, 64(4):18–23.

16. Prasertbun R, Mori H, Pintong AR, Sanyanusin S, Popruk S, Komalamisra C,Changbunjong T, Buddhirongawatr R, Sukthana Y, Mahittikorn A: Zoonotic potential of Enterocytozoon genotypes in humans and pigs in Thailand. Veterinary Parasitology 2017, 233:73–79.

17. Li W, Li Y, Li W, Yang J, Song M, Diao R, Jia H, Lu Y, Zheng J, Zhang X: Genotypes of Enterocytozoon bieneusi in livestock in China: high prevalence and zoonotic potential. Plos One 2014, 9(5):e97623.

18. Zhao W, Zhang W, Yang F, Cao J, Liu H, Yang D, Shen Y, Liu A: High prevalence of Enterocytozoon bieneusi in asymptomatic pigs and assessment of zoonotic risk at the genotype level. Applied & Environmental Microbiology 2014, 80(12):3699–3707.

19. Sulaiman IM, Fayer R, Lal AA, Trout JM, Iii FWS, Xiao L: Molecular Characterization of Microsporidia Indicates that Wild Mammals Harbor Host-Adapted Enterocytozoon spp. as well as Human-Pathogenic Enterocytozoon bieneusi. Applied & Environmental Microbiology 2003, 69(8):4495–4501.

20. Kumar S, Stecher G, Tamura K: MEGA7: Molecular Evolutionary Genetics Analysis Version 7.0 for Bigger Datasets. Molecular Biology & Evolution 2016, 33(7):1870.

21. Santín M, Fayer R: Enterocytozoon bieneusi genotype nomenclature based onthe internal transcribed spacer sequence: a consensus. Journal of Eukaryotic Microbiology 2009, 56(1):34–38.

22. Abe N, Kimata I: Molecular survey of Enterocytozoon bieneusi in a Japanese porcine population. Vector Borne & Zoonotic Diseases 2010, 10(4):425.

23. Piekarska J, Kicia M, Wesoå,Owska M, Kopacz Å, Gorczykowski M,Szczepankiewicz B, Kvã ÄM, Sak B: Zoonotic microsporidia in dogs and cats in Poland. Veterinary Parasitology 2017, 246:108–111.

24. Tavalla M, Mardanikateki M, Abdizadeh R, Soltani S, Saki J: Molecular diagnosis of potentially human pathogenic Enterocytozoon bieneusi and Encephalitozoon species in exotic birds in Southwestern Iran. J Infect Public Health 2017.

25. Deng L, Li W, Yu X, Gong C, Liu X, Zhong Z, Xie N, Lei S, Yu J, Fu H: First Report of the Human-Pathogenic Enterocytozoon bieneusi from Red-Bellied Tree Squirrels (Callosciurus erythraeus) in Sichuan, China. Plos One 2016, 11(9):e0163605.

26. Ke S, Li M, Wang X, Li J, Karim MR, Wang R, Zhang L, Jian F, Ning C: Molecular survey of Enterocytozoon bieneusi in sheep and goats in China. Parasites & Vectors 2016, 9(1):23.

27. Ding S, Huang W, Qin Q, Tang J, Liu H: Genotype Identification and Phylogenetic Analysis of Enterocytozoon Bieneusi Isolates from Stool Samples of Diarrheic Children. Journal of Parasitology 2018.

28. Tang C, Cai M, Wang L, Guo Y, Li N, Feng Y, Xiao L: Genetic diversity within dominant Enterocytozoon bieneusi genotypes in pre-weaned calves. Parasites & Vectors 2018, 11(1):170.

29. Wang L, Zhang H, Zhao X, Zhang L, Zhang G, Guo M, Liu L, Feng Y, Xiao L: Zoonotic Cryptosporidium species and Enterocytozoon bieneusi genotypes in HIV-positive patients on antiretroviral therapy. Journal of Clinical Microbiology 2013, 51(2):557.

30. Wei L, Lei D, Wu K, Huang X, Yuan S, Su H, Hu Y, Fu H, Zhong Z, Peng G: Presence of zoonotic Cryptosporidium scrofarum, Giardia duodenalis assemblage A and Enterocytozoon bieneusi genotypes in captive Eurasian wild boars (Sus scrofa) in China: potential for zoonotic transmission. Parasites & Vectors 2017, 10(1):10.

31. Li W, Tao W, Jiang Y, Diao R, Yang J, Xiao L: Genotypic distribution and phylogenetic characterization of Enterocytozoon bieneusi in diarrheic chickens and pigs in multiple cities, China: potential zoonotic transmission. Plos One 2014, 9(9):e108279–e108279.

32. Wan Q, Lin Y, Mao Y, Yang Y, Li Q, Zhang S, Jiang Y, Tao W, Li W: High Prevalence and Widespread Distribution of Zoonotic Enterocytozoon bieneusi Genotypes in Swine in Northeast China: Implications for Public Health. Journal of Eukaryotic Microbiology 2016, 63(2):162.

33. Li W, Diao R, Yang J, Xiao L, Lu Y, Li Y, Song M: High diversity of human-pathogenic Enterocytozoon bieneusi genotypes in swine in northeast China. Parasitology Research 2014, 113(3):1147.

34. Tian GR, Zhao GH, Du SZ, Hu XF, Wang HB, Zhang LX, Yu SK: First report of Enterocytozoon bieneusi from giant pandas (Ailuropoda melanoleuca) and red pandas (Ailurus fulgens) in China. In: Infect Genet Evol 34, 32-352015: 34, 32–35.

35. Wang L, Xiao L, Duan L, Ye J, Guo Y, Guo M, Liu L, Feng Y: Concurrent Infections of Giardia duodenalis, Enterocytozoon bieneusi, and Clostridium difficile in Children during a Cryptosporidiosis Outbreak in a Pediatric Hospital in China. PLoS Neglected Tropical Diseases,7,9(2013-9-12) 2013, 7(9):749–754.

36. Yang J, Song M, Wan Q, Li Y, Lu Y, Jiang Y, Tao W, Li W: Enterocytozoon bieneusi genotypes in children in Northeast China and assessment of risk of zoonotic transmission. Journal of Clinical Microbiology 2014, 52(12):4363.

37. Liu H, Jiang Z, Yuan Z, Yin J, Wang Z, Yu B, Zhou D, Shen Y, Cao J: Infection by and genotype characteristics of Enterocytozoon bieneusi in HIV/AIDS patients from Guangxi Zhuang autonomous region, China. Bmc Infectious Diseases 2017, 17(1):684.

38. Karim MR, Dong H, Li T, Yu F, Li D, Zhang L, Li J, Wang R, Li S, Li X: Predomination and new genotypes of Enterocytozoon bieneusi in captive nonhuman primates in zoos in China: high genetic diversity and zoonotic significance. Plos One 2015, 10(2):e0117991.

39. Karim MR, Wang R, Dong H, Zhang L, Li J, Zhang S, Rume FI, Qi M, Jian F, Sun M: Genetic Polymorphism and Zoonotic Potential of Enterocytozoon 18 bieneusi from Nonhuman Primates in China. Applied & Environmental Microbiology 2014, 80(6):1893.

40. Hu Y, Feng Y, Huang C, Xiao L: Occurrence, Source, and Human Infection Potential of Cryptosporidium and Enterocytozoon bieneusi in Drinking Source Water in Shanghai, China during a Pig Carcass Disposal Incident. Environmental Science & Technology 2014, 48(24):14219–14227.

41. Li N, Xiao L, Wang L, Zhao S, Zhao X, Duan L, Guo M, Liu L, Feng Y: Molecular surveillance of Cryptosporidium spp., Giardia duodenalis, and Enterocytozoon bieneusi by genotyping and subtyping parasites in wastewater. PLoS Neglected Tropical Diseases,6,9(2012-9-6) 2012, 6(9):e1809.

42. Ma J, Feng Y, Hu Y, Villegas EN, Xiao L: Human infective potential of Cryptosporidium spp., Giardia duodenalis and Enterocytozoon bieneusi in urban wastewater treatment plant effluents. Journal of Water & Health 2016, 14(3).

43. Qi M, Li J, Zhao A, Cui Z, Wei Z, Jing B, Zhang L: Host specificity of Enterocytozoon bieneusi genotypes in Bactrian camels (Camelus bactrianus) in China. Parasites & Vectors 2018, 11(1):219.

44. Zhao W, Zhang W, Yang Z, Liu A, Zhang L, Yang F, Wang R, Ling H: Genotyping of Enterocytozoon bieneusi in Farmed Blue Foxes (Alopex lagopus) and Raccoon Dogs (Nyctereutes procyonoides) in China. Plos One 2015, 10(11):e0143992.

45. Li W, Li Y, Song M, Lu Y, Yang J, Tao W, Jiang Y, Wan Q, Zhang S, Xiao L: Prevalence and genetic characteristics of Cryptosporidium, Enterocytozoon bieneusi and Giardia duodenalis in cats and dogs in Heilongjiang province, China. Veterinary Parasitology 2015, 208(3-4):125–134.

46. Karim MR, Dong H, Yu F, Jian F, Zhang L, Wang R, Zhang S, Rume FI, Ning C, Xiao L: Genetic diversity in Enterocytozoon bieneusi isolates from dogs and cats in China: host specificity and public health implications. Journal of Clinical Microbiology 2014, 52(9):3297–3302.

47. Li J, Luo N, Wang C, Meng Q, Cao J, Cui Z, Huang J, Wang R, Zhang L: 19 Occurrence, molecular characterization and predominant genotypes of Enterocytozoon bieneusi in dairy cattle in Henan and Ningxia, China. Parasites & Vectors 2016, 9(1):1–5.

48. Hu S, Liu Z, Yan F, Zhang Z, Zhang G, Zhang L, Jian F, Zhang S, Ning C, Wang R: Zoonotic and host-adapted genotypes of Cryptosporidium spp., Giardia duodenalis and Enterocytozoon bieneusi in dairy cattle in Hebei and Tianjin, China. Veterinary Parasitology 2017:68–73.

49. Meng Q, Bo J, Jian F, Wang R, Zhang S, Wang H, Ning C, Zhang L: Dominance of Enterocytozoon bieneusi genotype J in dairy calves in Xinjiang, Northwest China. Parasitology International 2017, 66(1):960–963.

50. Zhang Z, Huang J, Karim MR, Zhao J, Dong H, Ai W, Li F, Zhang L, Wang R: Zoonotic Enterocytozoon bieneusi genotypes in Pere David’s deer (Elaphurus davidianus) in Henan, China. Experimental Parasitology 2015, 155:46–48.

51. Huang J, Zhang Z, Yang Y, Wang R, Zhao J, Jian F, Ning C, Zhang L: New Genotypes of Enterocytozoon bieneusi Isolated from Sika Deer and Red Deer in China. Frontiers in Microbiology 2017, 8.

52. Qi M, Wang R, Wang H, Jian F, Li J, Zhao J, Dong H, Zhu H, Ning C, Zhang L: Enterocytozoon bieneusi Genotypes in Grazing Horses in China and Their Zoonotic Transmission Potential. Journal of Eukaryotic Microbiology 2016, 63(5):591–597.

